# Cellular heterogeneity of the LH receptor and its significance for cyclic GMP signaling in mouse preovulatory follicles

**DOI:** 10.1101/2020.02.06.937995

**Authors:** Valentina Baena, Corie M. Owen, Tracy F. Uliasz, Katie M. Lowther, Siu-Pok Yee, Mark Terasaki, Jeremy R. Egbert, Laurinda A. Jaffe

## Abstract

Meiotic arrest and resumption in mammalian oocytes are regulated by two opposing signaling proteins in the cells of the surrounding follicle: the guanylyl cyclase NPR2, and the luteinizing hormone receptor (LHR). NPR2 maintains a meiosis-inhibitory level of cyclic GMP (cGMP) until LHR signaling causes dephosphorylation of NPR2, reducing NPR2 activity, lowering cGMP to a level that releases meiotic arrest. However, the signaling pathway between LHR activation and NPR2 dephosphorylation remains incompletely understood, due in part to imprecise information about the cellular localization of these two proteins. To investigate their localization, we generated mouse lines in which HA epitope tags were added to the endogenous LHR and NPR2 proteins, and used immunofluorescence and immunogold microscopy to localize these proteins with high resolution. The results showed that the LHR protein is absent from the cumulus cells and inner mural granulosa cells, and is present in only 13-48% of the outer mural granulosa cells. In contrast, NPR2 is present throughout the follicle, and is more concentrated in the cumulus cells. Less than 20% of the NPR2 is in the same cells that express the LHR. These results suggest that to account for the LH-induced inactivation of NPR2, LHR-expressing cells send a signal that inactivates NPR2 in neighboring cells that do not express the LHR. An inhibitor of gap junction permeability attenuates the LH-induced cGMP decrease in the outer mural granulosa cells, consistent with this mechanism contributing to how NPR2 is inactivated in cells that do not express the LHR.

## Introduction

In mammalian preovulatory follicles, meiotic arrest is maintained by cyclic GMP (cGMP) that diffuses into the oocyte through gap junctions (1–3). The cGMP is generated in the granulosa cells of the follicle by the guanylyl cyclase natriuretic peptide receptor 2 (NPR2, also called guanylyl cyclase B) (4,5). cGMP then passes through the multilayer tissue to the oocyte by way of gap junctions formed by connexin 43 and connexin 37 (6–8). Regulation of meiotic arrest by cGMP generated by NPR2 appears to be conserved among mammals (1) including humans (9).

Mammalian oocytes resume meiosis in response to luteinizing hormone that activates a G-protein-coupled receptor (the luteinizing hormone / chorionic gonadotropin receptor, LHCGR, referred to here as LHR, since mice do not produce chorionic gonadotropin). LHR activation initiates signals that cause cGMP in the granulosa cells to decrease (1, 3). The decrease in cGMP to a meiosis-permissive level requires the dephosphorylation and inactivation of NPR2 (10–12). Phosphorylation and activation of the PDE5 cGMP phosphodiesterase, and other unknown mechanisms, also contribute to the LH-induced cGMP decrease (13). Due to diffusion through gap junctions, cGMP then decreases in the oocyte (6,7,14).

While the necessity of NPR2 inactivation for LH-induced meiotic resumption is established (12), the signaling pathway between the LHR and NPR2 dephosphorylation remains incompletely understood, due in part to imprecise information about the localization of these two proteins. In particular, it is unknown whether the approximately 50% decrease in NPR2 activity that occurs in response to LH signaling (10, 11) could result from inactivation of NPR2 solely in cells that express the LHR, or whether NPR2 in other cells must also be inactivated.

In previous studies, the localization of the LHR and NPR2 has been investigated by binding of radiolabeled mRNA probes and receptor ligands. These studies indicated that the LHR is located in the outer mural granulosa cell region of preovulatory follicles and in the theca and interstitial cells surrounding the follicle (15–20). mRNA encoding NPR2 is located in all regions of the granulosa compartment, but not in the oocyte (4, 21). NPR2 mRNA is more concentrated in the cumulus cell region (the granulosa cells directly surrounding the oocyte) and in the region of the mural granulosa closest to the antral space. However, these previous studies involved autoradiographic imaging and did not provide cellular level resolution, due to the scattering of the radiation emitted by the ~7-20 *μ*m thick histological sections, as well as the thickness of the overlying silver emulsion. In addition, mRNA localization does not provide direct information about protein levels. Antibodies that allow definitive immunolocalization of LHR and NPR2 within the follicle have not been developed, likely due to the difficulties of generating antibodies against G-protein coupled receptors and other transmembrane proteins (22), as well as the low expression levels of the LHR and NPR2.

To overcome this problem, we generated mice with a 9-amino acid hemagglutinin (HA) tag added to the extracellular N-termini of the endogenous LHR and NPR2 proteins. Because highly specific antibodies that recognize the HA peptide are available, this strategy has been used effectively to immunolocalize endogenous membrane proteins with low expression levels (23, 24). Using immunofluorescence and confocal microscopy, as well as immunogold and serial section electron microscopy, we investigated the localization of the LHR and NPR2 in preovulatory follicles, with much higher resolution than possible with previous methods. The distinct cellular localizations of these proteins indicate that LH-induced inactivation of NPR2 requires intercellular communication

## Materials and Methods

### Mice

CRISPR/Cas9 genome editing was used to generate two mouse lines in which a 9-amino acid HA epitope tag (YPYDVPDYA) was added to the extracellular N-termini of the endogenous LHR and NPR2 proteins; these modified proteins and mouse lines are referred to here as HA-LHR and HA-NPR2. The positions where the HA sequences were inserted (Fig. 1A,B) were the same as previously used to tag the LHR (25) and NPR2 (26,27) in cell lines.

**Figure 1.**
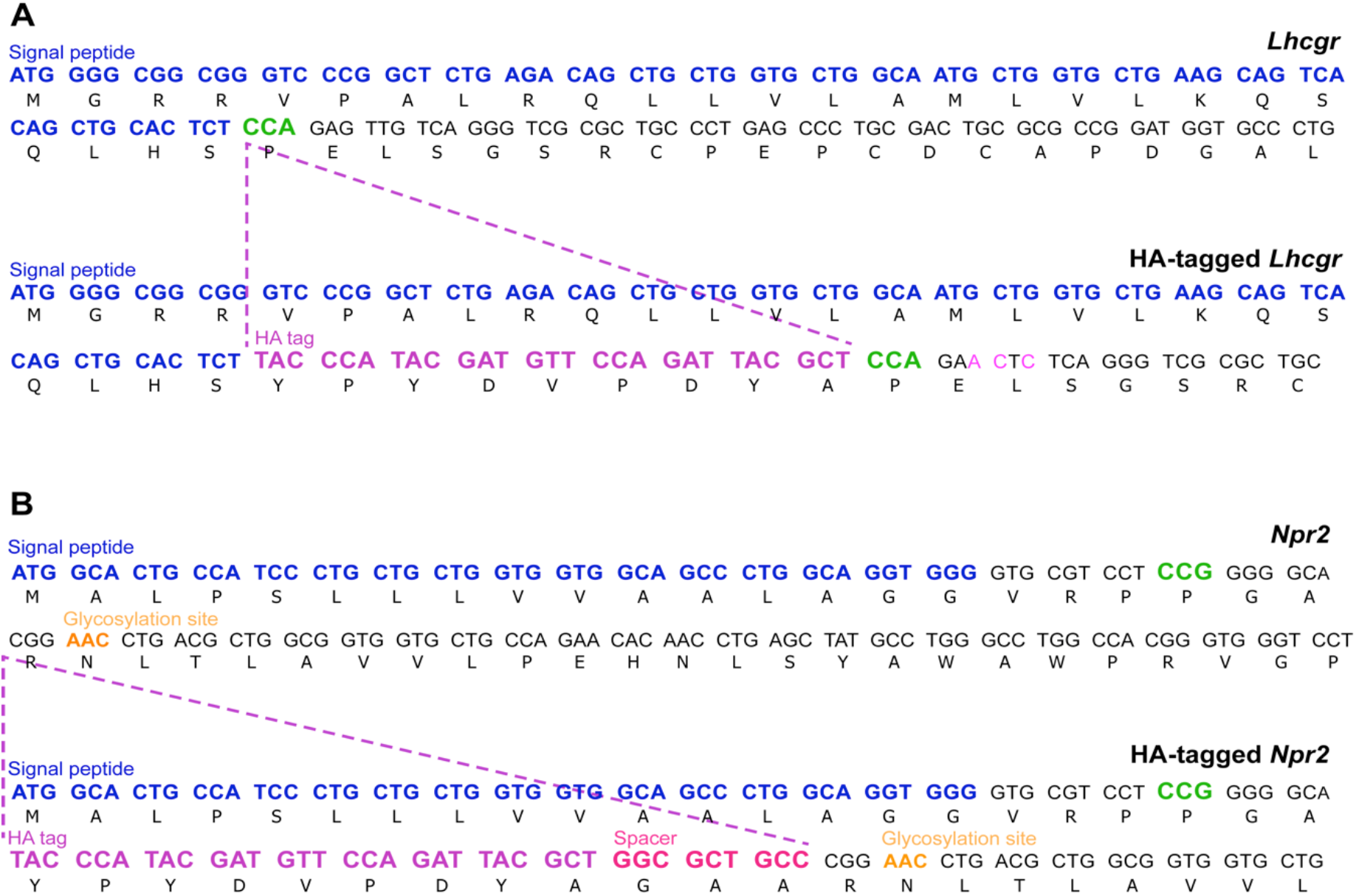
Generation of HA-LHR and HA-NPR2 mouse lines. (A) CRISPR-mediated gene editing was used to insert an HA tag (shown in purple) together with three silent point mutations (shown in pink) in the N-terminus of the *Lhcgr* coding sequence. The point mutations were located adjacent to the PAM site (shown in green) to eliminate re-cleavage of the HA-tagged *Lhcgr* allele. (B) An HA tag followed by a 3 amino acid spacer (shown in pink) was inserted into the *Npr2* coding sequence right after alanine 22 in the N-terminus. The spacer was used to separate the HA tag from an NPR2 glycosylation site (shown in orange). Signal peptides are shown in blue, and coding sequences in black.

Single guide RNAs (sgRNA) with sequences specific to the *Lhcgr* and *Npr2* genes, and single stranded DNA (ssDNA) donors for *Lhcgr* and *Npr2*, were obtained from Integrated DNA Technologies (Table 1). Cas9 protein was purchased from MilliporeSigma. Cas9 protein (50 ng/*μ*l) and sgRNA (25 ng/*μ*l) were incubated for 10 min at room temperature in injection buffer (10 mM Tris, pH 7.5, 0.1mM EDTA) and then mixed with ssDNA donor (40 ng/*μ*l).

**Table 1.**
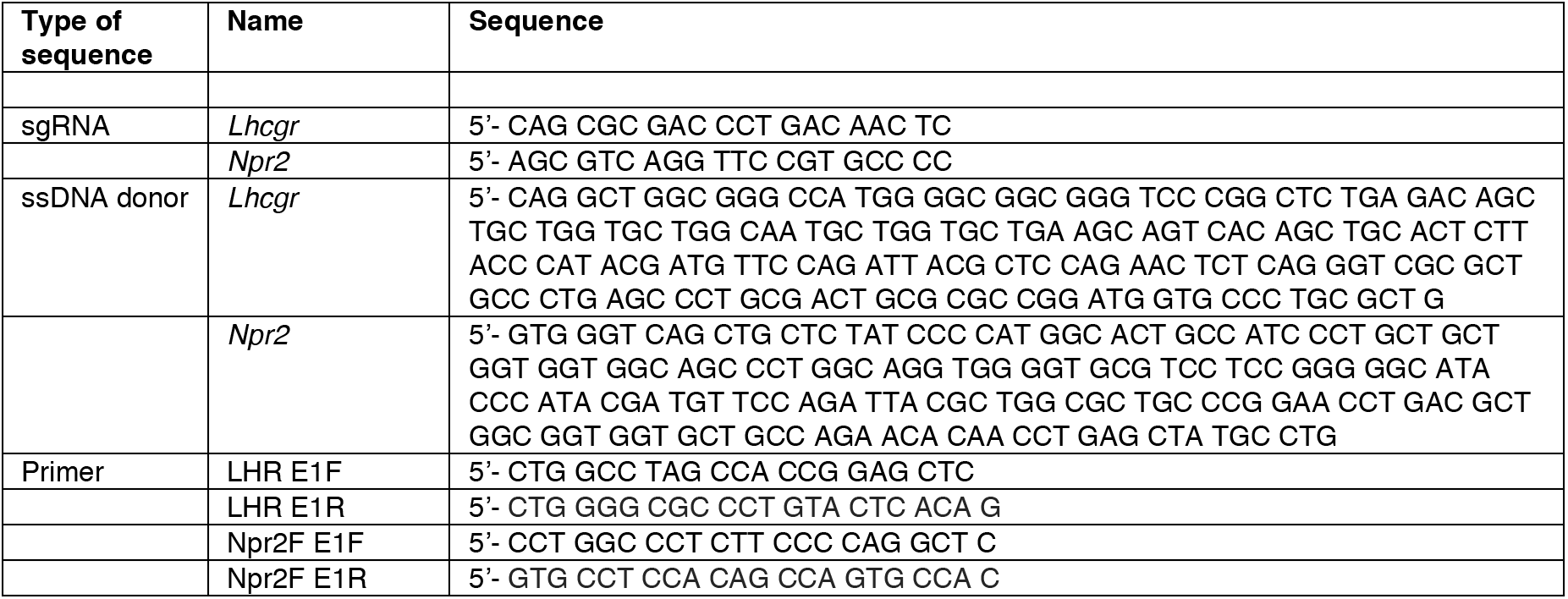
DNA sequences for generation and genotyping of HA-LHR and HA-NPR2 mice.

The Cas9/sgRNA ribonucleoprotein (RNP) complex and ssDNA donor mixture was microinjected into the pronuclei of C57BL/6J one-cell embryos. Injected embryos were transferred into CD1 foster females for subsequent development. Founder animals were initially identified by PCR using primer pairs (LHR E1F and LHR E1R, Table 1) to amplify fragments of 232 bp and 260 bp specific for *Lhcgr* and HA-tagged *Lhcgr*, respectively, and primer pairs (Npr2F E1F and Npr2F E1R, Table 1) to amplify fragments of 270 bp and 307 bp specific for *Npr2* and HA-tagged *Npr2*. Founder animals were confirmed by PCR followed by sequencing of the amplicon using the primer pairs as described above. Founders were then bred with wildtype C57BL/6J mice to expand the line, and the mice were maintained on a C57BL/6J background.

The HA-LHR and HA-NPR2 mouse lines were maintained by breeding of homozygotes, and all studies were performed using homozygous mice and wildtype controls. Fertility data were obtained by counting the cumulative number of pups produced over 4 months by each homozygous breeding pair, and by wild-type breeding pairs maintained in parallel.

An additional mouse line, expressing a sensor for cGMP (cGi500; 28), was provided by Robert Feil (University of Tübingen), and was maintained on a C57BL/6J background. Imaging studies using the cGi500 mice were performed using heterozygotes.

Mice were housed in a room with a 14 hour light/10 hour dark cycle (lights on at 0200 h, off at 1600 h). All experiments were conducted as approved by the University of Connecticut Health Center animal care committee.

### Ovaries, ovarian follicles, and gonadotropins

Except as indicated, ovaries were obtained by injecting 22-24 day old immature mice intraperitoneally with equine chorionic gonadotropin (eCG, 5 I.U.; National Hormone and Peptide Program, AFPSuperOv) to stimulate follicle growth to the preovulatory stage. 44 hours later, ovaries were dissected. To prepare samples for western blots, follicles ~470-540 *μ*m diameter were manually dissected from eCG-injected mice.

To obtain ovaries from adult mice on the day of proestrus, the stage of the estrous cycle was visually assessed by observation of the vaginal opening (29). Ovaries from mice that appeared to be at proestrus were collected at 1030 h to 1200 h, since serum LH levels begin to rise approximately one hour before the mouse room lights are turned off (30), and the lights in our mouse room were turned off at 1600 h. In this way, we obtained ovaries close to the time of the LH surge but avoided the possibility that the surge had already begun. The proestrous staging was confirmed by the presence of preovulatory follicles in ovary sections.

To test the effect of FSH on LHR protein expression, to determine the time course of nuclear envelope breakdown in response to LH, and to measure LHR mRNA by quantitative RT-PCR, follicles (~320-400 *μ*m diameter) were isolated from 23-26 day old mice that had not been injected with eCG. The isolated follicles were cultured for 24 hours with 1 nM FSH to stimulate LHR expression (6,31). Where indicated, the follicles were then incubated with 10 nM or 100 nM LH. Highly purified ovine FSH and LH were obtained from the National Hormone and Peptide Program (AFP7558C and LH-26, respectively). Nuclear envelope breakdown was scored by observation of intact follicles cultured on organotypic membranes (Millipore, Cork IR) such that the prophase arrested nucleus (germinal vesicle) could be seen within the follicle-enclosed oocyte (12).

### Western blots

Samples were prepared for SDS-PAGE by sonicating follicles in sample buffer with protease and phosphatase inhibitors (6). For all blots, 20 *μ*g of protein was loaded per lane. The protein content of preovulatory follicles from eCG-injected mice is ~6 *μ*g (Pierce BCA protein assay kit, 23225, Thermo Fisher Scientific). The protein content of follicles (~320-400 *μ*m diameter) from mice that were not injected with eCG, and which were treated with FSH in vitro, is ~3.5 *μ*g (6). Antibodies and stock concentrations are listed in Table 2. The primary antibody against the HA tag (32) was used at a dilution of 1:1000. The secondary antibody (33) was used at a dilution of 1:20,000. Blots were developed with a WesternBright Sirius Chemiluminescent Detection kit (Advansta, K-12043).

**Table 2.** Antibodies

| Name | Source | Stock concentration |
| --- | --- | --- |
| rabbit monoclonal antibody against HA tag; RRID: AB_1549585 | Cell Signaling Technology, 3724, lot 9 | 67 μg/ml |
| goat anti-rabbit IgG (H+L), HRP conjugate; RRID: AB_10719218 | Advansta, R-05072-500 | 1 mg/ml |
| goat anti-rabbit IgG 488-Alexa Fluor secondary antibody; RRID: AB_2576217 | Thermo Fisher Scientific, Molecular Probes, A-11034 | 2 mg/ml |
| goat anti-rabbit IgG Alexa Fluor 488 - FluoroNanogold secondary antibody; RRID: AB_2819209 | Nanoprobes, 7203 | 80 μg/ml |

### Quantitative RT-PCR

Total RNA was purified from isolated follicles that had been cultured for 24 hours with FSH as described above. RNA was isolated using the RNeasy Micro kit (Qiagen) and reverse transcribed using the SuperScript III kit (Invitrogen) with random primers according to manufacturer’s instructions. Forward and reverse primers (Table 3) were validated by performing a standard curve with cDNA to determine the linear range of each set of primers. Quantitative RT-PCR was performed using the iTaq SYBR Green Supermix (Bio-Rad) and a CFX Connect Detection System (Bio-Rad). Assays were performed in duplicate. The cycle threshold method was used to calculate relative expression levels after normalization to TATA binding protein *(Tbp)* levels

**Table 3.**
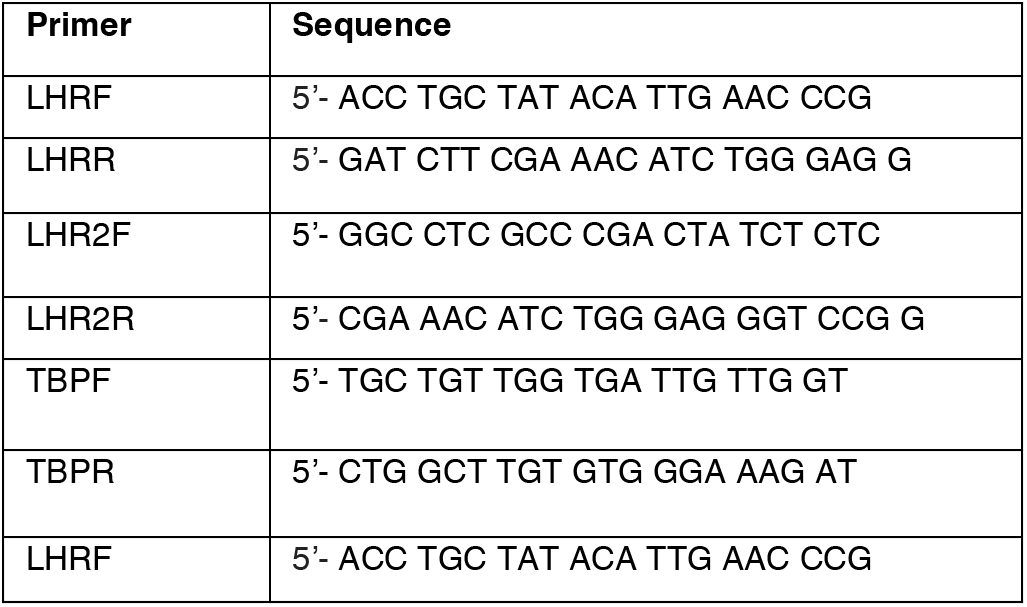
Primers for RT-PCR quantitation of mRNA encoding LHR

### Immunofluorescence labeling of ovary sections

Ovaries were fixed in 4% paraformaldehyde (a 1:1 mixture of 8% paraformaldehyde, Electron Microscopy Sciences (EMS) 157-8, and PBS) for 4 hours at room temperature, then rinsed 3 times in PBS (5 minutes each). The fixed ovaries were transferred to 15% sucrose at 4°C until they sank, transferred to 30% sucrose at 4°C until they sank, then embedded in OCT medium (EMS 62550-01) and frozen in a slurry of dry ice and ethanol. Ten or 20 *μ*m thick cryostat sections were collected on slides and rinsed 3 times with PBS. Sections were blocked with 5% normal goat serum and 1% BSA in PBS, then labeled with the HA-tag antibody (32) listed in Table 2 (1:500 dilution in the blocking solution) for 1 hour at room temperature. The sections were washed with PBS several times, labeled with a goat anti-rabbit 488-Alexa Fluor secondary antibody (34) (see Table 2, 1:500 dilution) for 1 hour at room temperature, then washed several times in PBS. Some sections were additionally labeled with 4′,6-diamidino-2-phenylindole (DAPI, Sigma D9542, 0.5 *μ*g/ml in PBS, for 15 min at room temperature, followed by 5 × 3 min washes in PBS) to label nuclei. Labeled sections were mounted with Shandon Immu-Mount (Thermo Fisher Scientific 9990402) and a #1 coverslip.

### RNAscope labeling of ovary sections

RNAscope was performed using a RNAscope Fluorescent Multiplex Reagent kit (Advanced Cell Diagnostics, 320850) and a probe designed for *Lhcgr* (Advanced Cell Diagnostics, 408171). Ovaries from eCG-injected mice were obtained as described above and frozen in OCT without fixation. RNAscope was performed according to the kit protocol. Briefly, 10 *μ*m thick cryosections were fixed in 4% paraformaldehyde for 15 minutes at 4°C, then dehydrated in 50%, 70%, 100%, and 100% ethanol for 5 minutes each. Slides were dried, treated with Protease IV for 30 minutes at 40°C, then washed twice with PBS. The probe was applied to the slide for 2 hours in a humidified chamber at 40°C followed by the 4 amplification steps using amplification reagent B (orange fluorophore). Slides were washed, treated with DAPI, and mounted with Shandon Immu-Mount.

### Confocal microscopy and analysis

Ovary sections were imaged with a confocal microscope (Pascal or LSM800, Carl Zeiss Microscopy) and images were saved as 12-bit or 16-bit files. For analysis of protein localization, the outer and inner regions of the mural granulosa layer were defined as follows: The width of the mural granulosa layer was measured from the basal lamina to the antrum at 8 radial points in the follicle, and the halfway point was marked. The points were connected to mark the boundary between the inner and outer mural cells.

To determine the proportion of outer mural granulosa cells that expressed the LHR, 10 *μ*m sections that had been co-stained with DAPI were used to count cells with the Cell Counter Tool in Fiji (35). DAPI-stained nuclei that were surrounded by HA labeling were counted as LHR-positive cells. DAPI-stained nuclei without HA labeling were counted as LHR-negative cells. Cell counts were made from a series of 10 optical sections taken at 1 *μ*m intervals and having an approximate optical section thickness of ~1.5 *μ*m. A similar method was used to determine the fraction of LHR-expressing cells in the layer of cells with cell bodies directly adjacent to the basal lamina.

To determine the relative expression levels of HA-NPR2 in each region of the follicle, z-stacks of confocal images were collected from 20 *μ*m thick sections located at the level of the oocyte. Intensity measurements were made from the central optical section, using Image J. Background intensities for each region were determined by performing similar measurements using wildtype follicles. The average background fluorescence intensity was subtracted from the intensities measured from HA-NPR2 follicles. Background-subtracted intensity values for each region of each HA-NPR2 follicle were normalized to the intensity of the cumulus cells.

### Immunogold labeling of vibratome slices of ovary

Ovaries were fixed in 4% paraformaldehyde (prepared as described above for immunofluorescence) overnight at 4°C, then rinsed 3 times in PBS (5 minutes each) and embedded in 4% low gelling temperature agarose in PBS (Sigma-Aldrich A0701). 55 *μ*m slices of the ovaries were cut with a vibratome (Leica VT 1000 S) and were collected and further processed in glass depression wells. After a few washes with PBS, the slices were permeabilized with Triton X-100 (Thermo Fisher Scientific, 28314) in PBS for 20 minutes with gentle shaking. 0.1% Triton X-100 was used to obtain optimal preservation for thin sectioning, and 0.3% Triton X-100 was used to achieve higher antigen labeling for thick sectioning.

All subsequent steps were carried out with gentle shaking at room temperature unless specified otherwise. Slices were rinsed once with PBS, treated with 5% normal goat serum, 1% BSA, and 0.1% Triton X-100 in PBS for 30 minutes to block non-specific binding, then labeled with the HA-tag antibody (32) listed in Table 2. The antibody was used at a 1:100 dilution (for thin sectioning) or 1:300 (for thick sectioning), in the blocking solution overnight. Slices were then washed with PBS 5 times for 5 minutes each and then treated with the blocking solution for 30 minutes. After blocking, slices were incubated in a secondary antibody conjugated to Alexa Fluor 488 and nanogold (36) (see Table 2), at a dilution of 1:100 (for thin sectioning) or 1:200 (for thick sectioning). The secondary antibody was diluted in the blocking solution but without Triton X-100, and was applied for 4 hours in the dark. Slices were then washed with PBS 5 times for 5 minutes each, placed in a drop of PBS on a glass slide, and the fluorescence was checked using an epifluorescence microscope.

Slices were then fixed with 1% glutaraldehyde diluted in PBS from an 8% aqueous solution (EMS 16020) for 10 minutes, followed by 3 PBS washes for 3 minutes each, then 5 washes with 0.1 M Tris buffer for 3 minutes each. The nanogold was enhanced with a gold enhancement kit (GoldEnhance EM Plus, Nanoprobes 2114) for 9 minutes following the manufacturer’s instructions. The enhancement was stopped by washing the slices with 0.1 M Tris buffer 5 times for 2 minutes each, followed by 5 washes with PBS for 2 minutes each. The slices were then transferred to capped glass vials and postfixed with 1% osmium tetroxide diluted in PBS from a 4% aqueous solution (EMS 19170) for 30 minutes on a rotator. The slices were washed with 5 rinses of Milli-Q water for 2 minutes each, then placed in 1% uranyl acetate in water (EMS 22400) and stored at 4°C overnight. The next day, they were washed with 5 rinses of Milli-Q water for 2 minutes each.

The slices were dehydrated in sequential ethanol solutions, infiltrated with Epon resin (Embed-812 kit, EMS 14121), and flat-embedded in the same resin between two pieces of ACLAR embedding film with a 50 *μ*m thickness (EMS 50426) using 2 pieces of Parafilm as spacers between them and a glass slide on top to provide weight to flatten the slices. The resin was polymerized in a 60°C oven for at least 20 hours, and the embedded slices were then attached to blank Epon blocks using superglue.

### Sectioning and imaging for electron microscopy, and analysis of images

Trimming, sectioning, imaging, and image alignment were done as previously described (37,38) with the following modifications: For thin sectioning of the samples that were treated to enhance the preservation of structures, serial sections were cut at a thickness of 65 nm. For thick sectioning of the samples that were treated to enhance antigen labeling, the serial sections were cut at 500 nm.

Reconstructions were made using the TrakEM2 module (39) from FIJI by manually segmenting cell outlines or the basal lamina on the serial sections using arealists. The surface areas of cell contacts with the basal lamina were estimated from the Upper Bound smoothened measurement of the arealists.

### Live confocal imaging of isolated follicles to measure cGMP levels

Measurements of cGMP were made using mice expressing the cGi500 Förster resonance energy transfer (FRET) sensor (28, 40). Methods were as previously described (13, 14), except that LH was used at a concentration of 10 nM instead of 300 nM. Carbenoxolone (CBX) was obtained from MilliporeSigma (C4790).

### Fluorescence recovery after photobleaching

Measurements were performed and analyzed as previously described (14).

### Statistics

All analyses were conducted using Prism 6 (GraphPad Software, La Jolla, CA). Details are given in the figure legends.

## Results

### Validation of mice with HA-tagged LHR and NPR2

To determine whether mice in which an HA epitope tag was added to the N-terminus of the endogenous LHR produced a protein of the expected size and with the expected developmental profile, proteins from isolated preovulatory follicles, obtained from eCG-stimulated mice, were examined by western blotting. As predicted from the HA-LHR sequence, an HA-positive band was seen at ~80 M_r_ (Fig. 2A). Some lower molecular weight bands were also present, presumably representing breakdown products or splice variants. The ~80 M_r_ and lower molecular weight bands were absent in follicles from wildtype mice, although one non-specific band at ~116 M_r_ was seen in both HA-LHR and wildtype follicles (Fig. 2A,B). Further confirming the appropriate expression of the HA-LHR protein, the ~80 M_r_ and lower molecular weight HA-positive bands were absent in follicles that had not been exposed to follicle-stimulating hormone (FSH), a hormone that stimulates expression of the LHR (15,16,41) (Fig. 2B).

**Figure 2.**
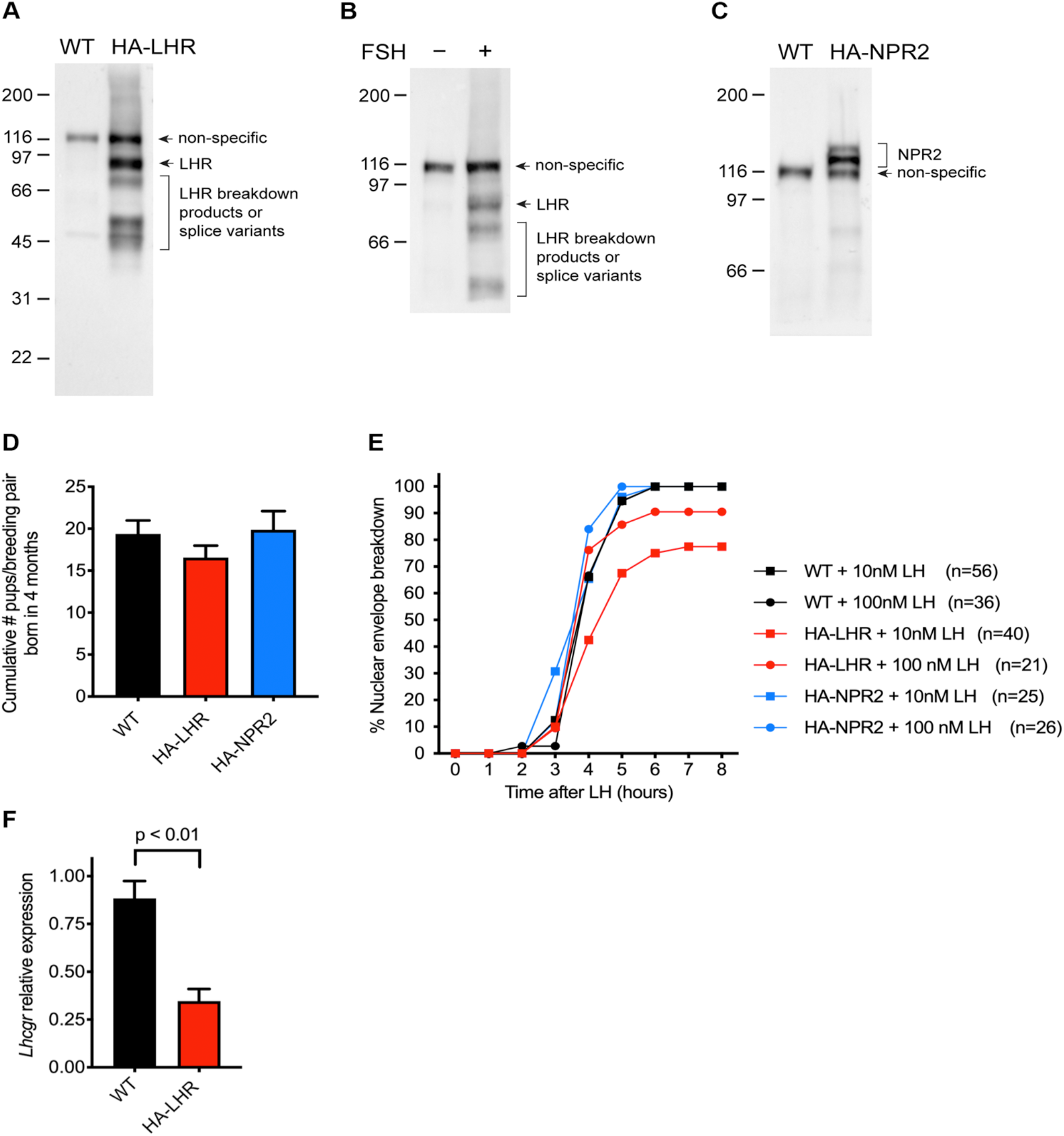
Validation of the size, expression, and biological activity of the HA-LHR and HA-NPR2 proteins in the genetically modified mice. (A-C) Western blots of preovulatory follicles, probed with an HA tag antibody. (A) Follicles from HA-LHR and wildtype (WT) mice. (B) Follicles from HA-LHR mice, with and without a 24 hour incubation of the follicles with FSH. (C) Follicles from HA-NPR2 and WT mice. (D) No obvious defect in fertility of HA-LHR and HA-NPR2 mice, as judged by the number of pups produced in 4 months by homozygous breeding pairs, compared with wild-type pairs (mean ± SEM; 9 pairs for WT, 14 pairs for HA-LHR, and 8 pairs for HA-NPR2). Data were analyzed by one-way ANOVA with the Holm-Sidak correction for multiple comparisons. No differences in the cumulative number of pups per breeding pair were observed (p > 0.05). (E) Time course of nuclear envelope breakdown in response to 10 or 100 nM LH, comparing HA-LHR, HA-NPR2, and WT follicles. Nuclear envelope breakdown was scored by observation of live follicles (6). The results were obtained from 5 independent experiments, 3 of which included parallel tests of HA-LHR and wildtype follicles incubated with 10 nM LH. n values indicate the total number of follicles tested for each condition. (F) Relative expression of *Lhcgr* mRNA in pools of 4-15 follicles cultured for 24 hours with FSH from WT and HA-LHR mice (n = 5 mice for each genotype). Total RNA was isolated and reverse transcribed, and levels of *Lhcgr* were measured using qRT-PCR with primers LHRF and LHRR (Table 3) and normalized to levels of *Tbp*. Data are reported as mean fold change ± SEM and were analyzed by an unpaired t test. Similar results were obtained by analysis of the same samples using a different pair of primers, LHR2F and LHR2R in Table 3.

Follicles from mice in which an HA epitope tag was added to the N-terminus of the endogenous NPR2 also showed expression of a protein of the appropriate molecular weight, at ~120 and ~130 M_r_ (Fig. 2C), as expected based on its known glycosylation (11). As with the HA-LHR mice, a non-specific band was present at ~116 M_r_. Ideally, different epitope tags would have been used for the two lines, to allow colocalization of the two proteins. We also made mice expressing NPR2 with a FLAG tag (DYKDDDDK; 42) or a PA tag (GVAMPGAEDDVV; 43) in the same position as described for the HA tag. However, in contrast to the HA-tagged mice described here, follicles from mice with FLAG-tagged or PA-tagged NPR2 did not show specific labeling with western blots and immunofluorescence. Thus, of the three tags that we compared, only HA was useful with the antibodies that we tested, probably due to low expression levels of NPR2.

HA-LHR and HA-NPR2 homozygous breeding pairs produced approximately the same number of pups as wildtype mice, indicating no obvious defect in their fertility (Fig. 2D). To further test whether these mice showed normal ovarian function, we examined the response of isolated follicles to LH. Oocytes within HA-LHR- and HA-NPR2-expressing follicles resumed meiosis, as indicated by nuclear envelope breakdown (Fig. 2E). For HA-LHR-expressing follicles, the response to 100 nM LH showed kinetics similar to wildtype, but the response to 10 nM LH was slightly slower than in wildtype and occurred in only about 75% of the follicles. This slight attenuation could potentially be due to an effect of the HA tag on the level of LHR mRNA, or on the binding affinity of the receptor. Quantitative PCR analysis indicated that the amount of LHR-encoding mRNA in follicles from HA-LHR-expressing mice was ~40% of that from wildtype mice (Fig. 2F). However, these differences were not sufficient to be obviously detrimental to fertility.

### Localization of the LHR in a subset of the outer mural granulosa cells

To investigate the localization of the LHR, we examined frozen sections of formaldehyde-fixed ovaries from HA-LHR-expressing mice; sections were labeled with an HA-tag antibody (32). Confirming previous mRNA and ligand binding studies (15–20), the LHR was localized in the outer layers of mural granulosa cells of preovulatory follicles, but not in the inner mural cells, cumulus cells, or oocyte (Fig. 3A,C). HA-antibody labeling was not seen in the mural granulosa cells of wildtype follicles (Fig. 3B,D), or in the granulosa cells of follicles that had not grown to the preovulatory stage (Fig. 3A). In ovaries of HA-LHR-expressing mice, a variable degree of HA-antibody labeling was also detected in some of the theca and interstitial cells that surround the follicle (see examples in figures 3 and 4).

**Figure 3.**
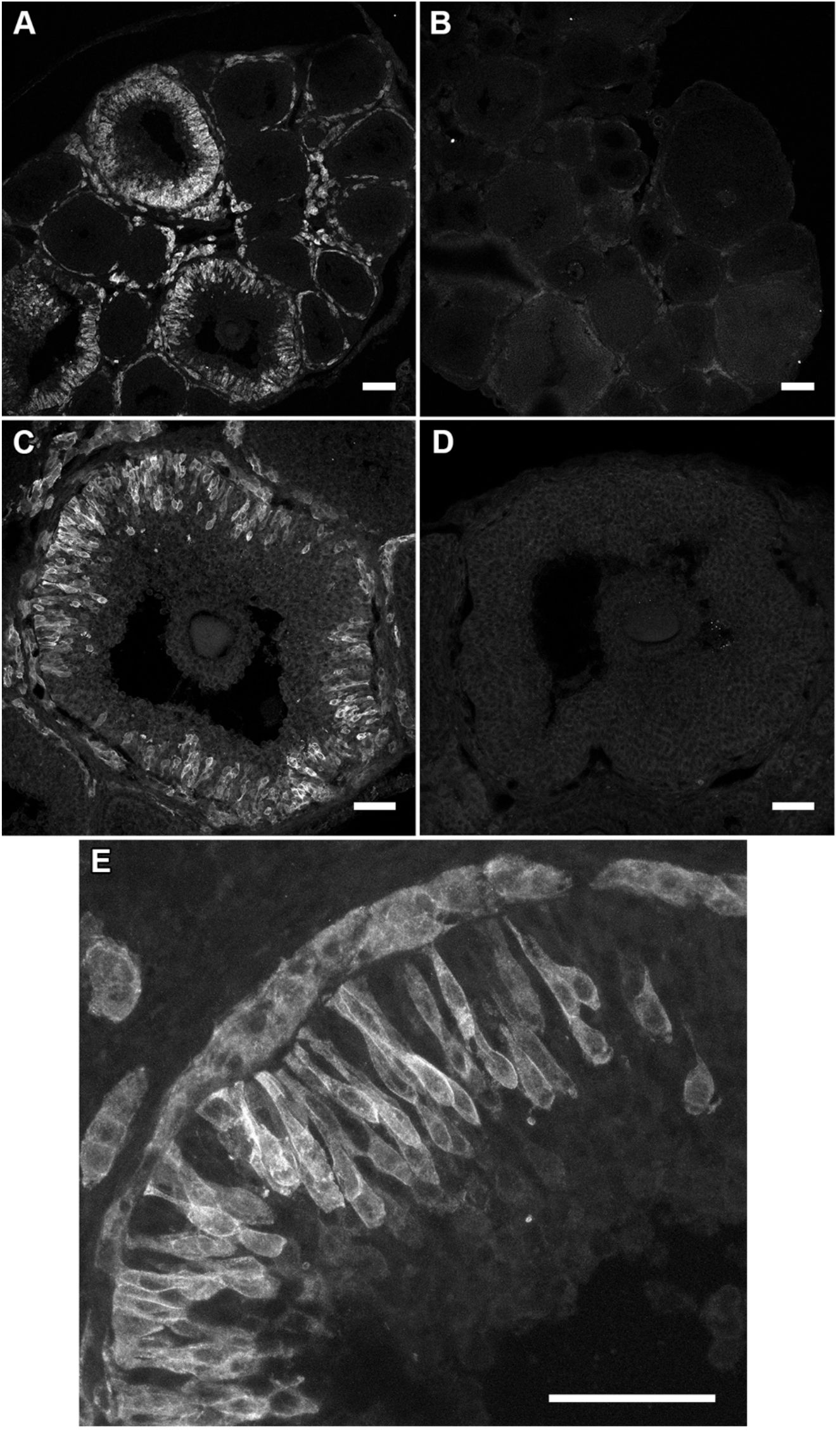
Cellular heterogeneity of the LHR in preovulatory follicles. Ovaries were collected from immature HA-LHR (A,C,E) or wildtype (B,D) mice that were injected 44 hours previously with eCG. All sections were labeled with an HA tag antibody. HA-LHR and wildtype sections were labeled and imaged using identical conditions. A and B show single optical sections. C-E show maximum projections of 11 optical sections imaged at 1 *μ*m intervals. Scale bars indicate 100 *μ*m for A and B, 50 *μ*m for C-E.

**Figure 4.**
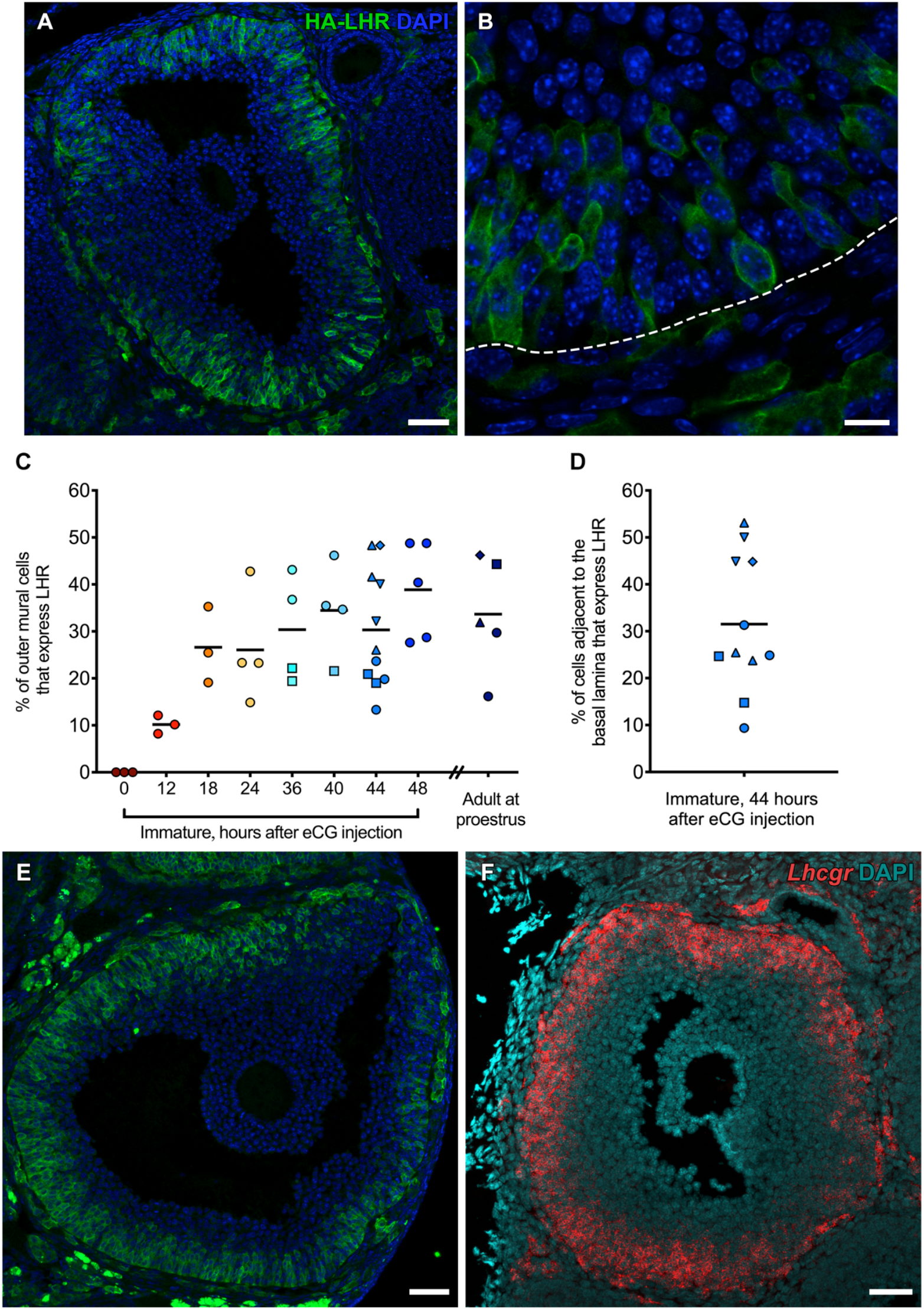
Quantitation of the fraction of outer mural granulosa cells that express the LHR, in immature mice after eCG injection and in adult mice at proestrus. (A) Preovulatory follicle from an ovary of an immature HA-LHR mouse that was injected 44 hours previously with eCG. HA-LHR in green, DAPI-labeled nuclei in blue. Maximum projection of 10 confocal images taken at 1 *μ*m intervals. (B) High magnification image of a single section from a region of the follicle in A. The dotted line indicates the basal lamina. Video 1 shows the stack of confocal images through the 10 *μ*m physical section. (C) Percentage of outer mural granulosa cells that express the LHR in immature mice at various times after injection of eCG, and in adult mice at proestrus. For each time point, different symbols indicate different mice, and the horizontal bar indicates the mean value. (D) Percentage of cells with cell bodies directly adjacent to the basal lamina that express the LHR, in immature mice at 44 hours after injection of eCG. (E) Preovulatory follicle in the ovary of an adult mouse at proestrus. Maximum projection of 10 confocal images taken at 1 *μ*m intervals. (F) Localization of the mRNA encoding the LHR in a preovulatory follicle from an ovary of an immature wildtype mouse that was injected 44 hours previously with eCG. mRNA in red, DAPI-labeled nuclei in blue. Representative of results from 11 follicles from 3 mice. Scale bars indicate 50 *μ*m for A, E, and F; 10 *μ*m for B.

Many of the LHR-expressing mural granulosa cells were flask-shaped, with long projections extending to the basal lamina from cell bodies several layers deep (Fig. 3E), comprising a pseudostratified epithelium (37, 44). Most of the LHR protein was located at the cell surface, although some appeared to be intracellular. Strikingly, only some of the cells in the outer half of the mural granulosa cell region of preovulatory follicles expressed the LHR (Fig. 3E and other examples in figures 3-5). Some areas of the outer mural region showed intense labeling, while others showed none. On a cellular level, some cells were labeled, while some adjacent cells were not.

To quantify the fraction of cells in the outer half of the mural granulosa region that expressed the LHR, we labeled cell nuclei with DAPI and counted the total number of cells in a 10 *μ*m thick equatorial section of an HA-LHR follicle (see Materials and Methods, Fig. 4A,B, and video 1). We also counted the number of these cells that expressed the LHR from images like that shown in Fig. 4B and video 1. In 13 preovulatory follicles from four mice injected 44 hours previously with the FSH receptor agonist eCG, LHR expression was seen in less than half of the outer mural granulosa cells, with percentages ranging from 13% to 48% (Fig. 4C). Similar results were obtained from mice injected with eCG 24, 36, 40 or 48 hours previously, although less labeling was seen at 12 hours after eCG injection and no labeling was seen in ovaries from mice without eCG injection (Fig. 4C). Analysis of the subpopulation of mural cells whose cell bodies are in direct contact with the basal lamina also showed that only 9-53% of these cells expressed the LHR (Fig. 4D)

The results described above were obtained using ovaries from immature (24-26 day old) mice. Although eggs obtained from such mice are capable of normal development (45), we also examined LHR expression in ovaries from adult mice on the day of proestrus. In these ovaries as well, only 16 to 46% of the outer mural granulosa cells expressed the LHR (Fig. 4C,E).

To test whether the HA tag might be causing the heterogenous expression pattern, we examined the localization of mRNA encoding the LHR in wildtype follicles. As seen for the LHR protein in HA-LHR follicles, the LHR mRNA in wildtype follicles was not uniformly expressed within the outer mural granulosa region (Fig. 4F).

### Localization of LHR by immunogold labeling and serial section electron microscopy

To compare the morphology of LHR-expressing and non-expressing cells, and to investigate their intercellular contacts, we used immunogold labeling and serial section electron microscopy to reconstruct the shapes of both cell types. 55 *μ*m thick vibratome slices of formaldehyde-fixed ovaries were permeabilized with Triton X-100, and the HA epitope was labeled with nanogold prior to embedding and sectioning. The gold label appeared black as viewed with transmitted light, and the labeling pattern confirmed that the LHR protein is distributed in a subset of the outer mural granulosa cells of preovulatory follicles, and in some cells outside of the basal lamina (Fig. 5A).

**Figure 5.**
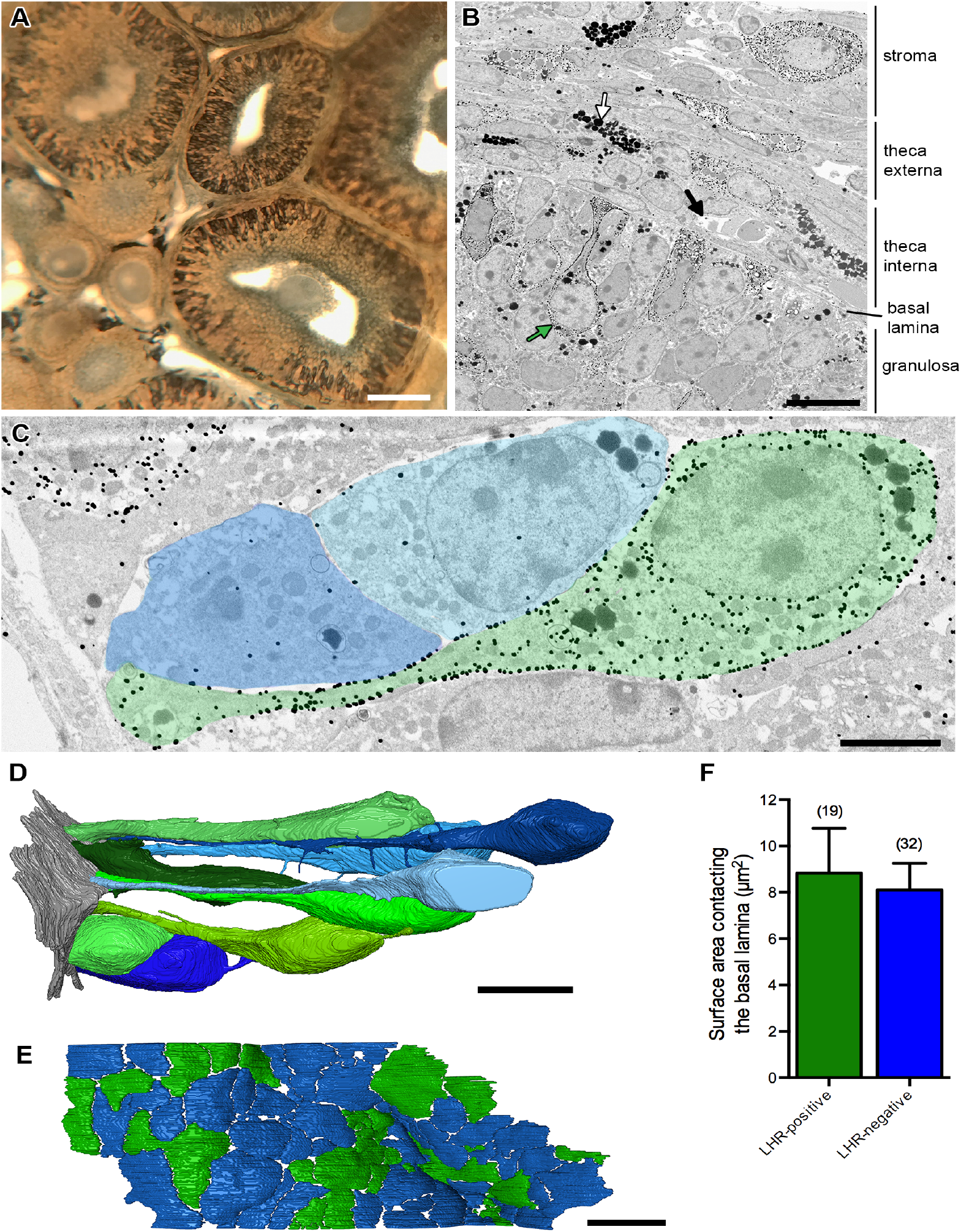
Electron microscopic imaging of the shapes and basal lamina contacts of mural granulosa cells that do or do not express the LHR. (A) Light microscope view of a 55 *μ*m thick vibratome slice of an ovary from an immature HA-LHR mouse that was injected 44 hours previously with eCG. The slice was permeabilized with 0.1% Triton X100, labeled with an HA tag antibody and immunogold, and flat-embedded in Epon resin. The black staining is the enhanced gold (LHR label). The diffuse gray staining around the antrum and in the oocyte is background labeling. (B) A 500 nm section of the vibratome slice, imaged by scanning electron microscopy. The small black dots indicate gold-labeling of the LHR (green arrow). The large black dots indicate lipid drops (white arrow) and do not indicate gold labeling. The black arrow points to a blood vessel adjacent to the basal lamina. (C) A 65 nm section from a series of 225 sections. LHR-expressing cells (green) were identified by the presence of gold labeling in the plasma membrane, nuclear envelope, and cytoplasm, throughout several serial sections. Two non-LHR-expressing cells, identified by the lack of gold labeling, are shown in blue. Basal lamina contact sites are visible on the left side of the figure. (D) Reconstruction of 9 cells traced from serial sections, with LHR-expressing cells shown in shades of green and non-LHR-expressing cells shown in shades of blue. The basal lamina surface is shown in gray on the left. Video 3 shows this reconstruction from multiple angles. (E) Reconstruction showing contact sites with the basal lamina for 71 cells. Contacts with LHR-expressing cells are shown in green and contacts with non-LHR-expressing cells are shown in blue. F) Average surface area of the contact site made by LHR-expressing cells and non LHR-expressing cells with the basal lamina. Bars show mean ± SEM. Means were compared using an unpaired t-test; p > 0.05. Numbers indicate the number of cell contacts measured for each group. Scale bars indicate 100 *μ*m for A, 10 *μ*m for B, 3 *μ*m for C, 10 *μ*m for D, and 5 *μ*m for E.

500 nm sections of these slices showed gold particles bound to some of the outer mural granulosa cells as well as to some of the cells in the theca interna, theca externa, and stroma layers (Fig. 5B). Multiple cell types with functions that are only partly known are present within these layers (46).

Analysis of 225 serial sections, each 65 nm thick, allowed precise definition of all of the LHR-expressing and non-LHR-expressing cells in a 15 *μ*m thick volume (Fig. 5C-E, video 2). The thin section images showed that the LHR protein was most concentrated in the plasma membrane, but some was in the nuclear envelope and cytoplasm (Fig. 5C, video 2). Tracing of individual cells through multiple serial sections showed that almost all of the LHR-expressing cells contacted the basal lamina, either at the cell body or by way of a long projection (Fig. 5C,D). The rare LHR-expressing cells that did not appear to contact the basal lamina were located near the middle of the granulosa layer. Non-LHR-expressing cells had similar shapes and also sent projections to the basal lamina, making similar contact sites (Fig. 5D-F). Thus, basal lamina contact is not sufficient for LHR expression. There was no obvious relationship between proximity to blood capillaries near the basal lamina (see Fig. 5B) and expression of the LHR in adjacent granulosa cells.

Serial section reconstruction also identified gap junctions (Fig. 6 and video 3) and fine cellular processes (Fig. 5D) connecting HA-LHR-expressing and non-expressing mural granulosa cells. As will be described below, these gap junctions function in signaling between the LH receptor and the NPR2 guanylyl cyclase in adjacent cells.

**Figure 6.**
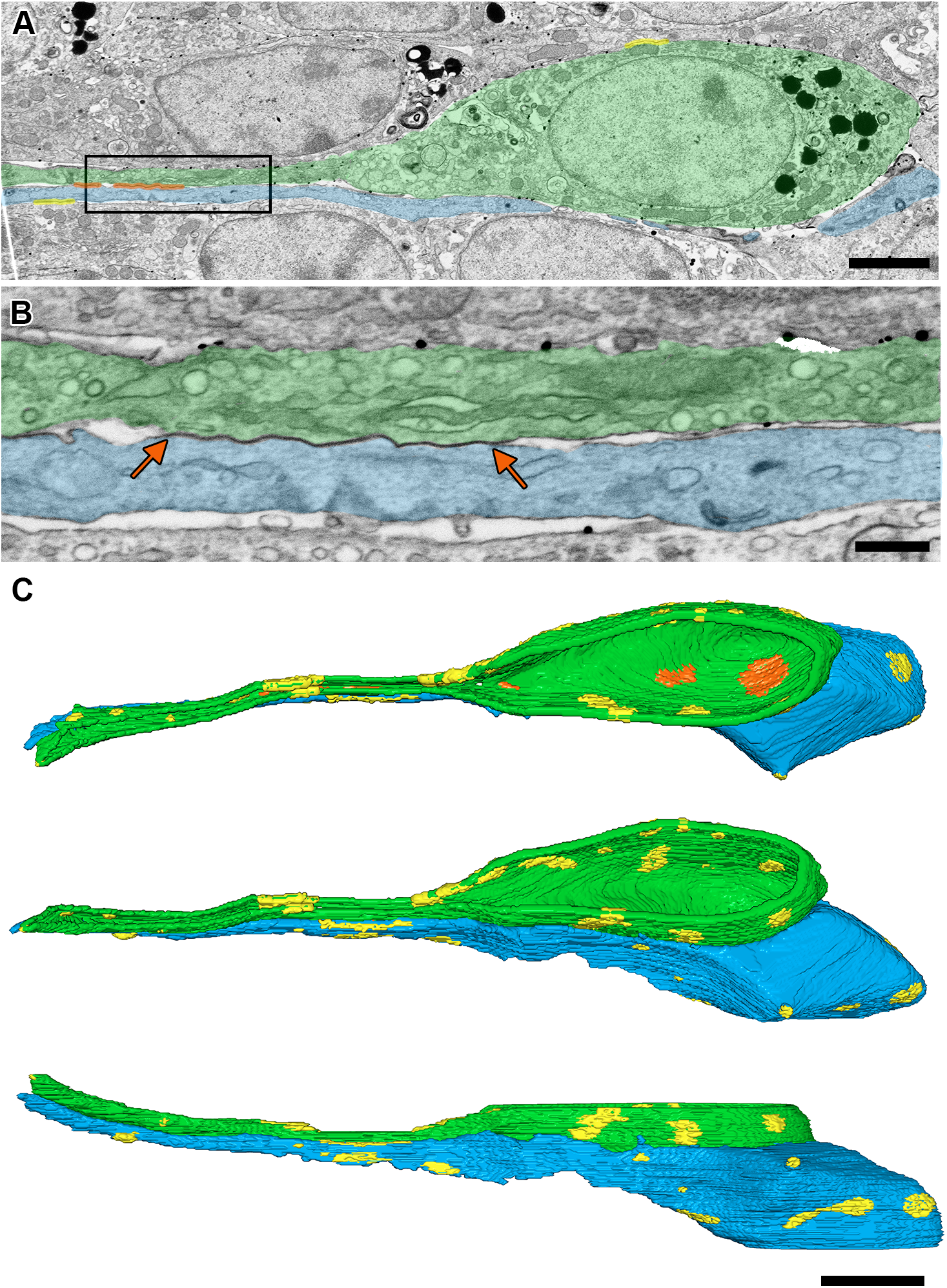
Gap junctions between LHR-expressing and non-LHR-expressing cells. (A) Gap junction contacts of an LHR-expressing cell (green) and a non-LHR-expressing cell (blue) are highlighted in orange. Yellow indicates gap junctions with other cells. The box in A indicates the area enlarged in B. (B) A gap junction (orange arrows) between the 2 cells highlighted in A. (C) Reconstruction of the 2 cells in A, showing 3 angles of view from video 4. Color coding of gap junctions as for A. Scale bars indicate 3 *μ*m for A, 500 nm for B, and 5 *μ*m for C.

### Localization of NPR2 throughout the granulosa cell compartment, with a higher concentration in the cumulus region

To investigate the distribution of NPR2 protein in the follicle, we examined frozen sections of formaldehyde-fixed ovaries from HA-NPR2-expressing mice by labelling sections with an HA-tag antibody (32). Consistent with previous reports of NPR2 mRNA localization (4,21), NPR2 protein was present throughout the granulosa region of the follicle, absent in the oocyte, and most concentrated in the cumulus cells (Fig. 7A; compare with the wildtype follicle in Fig. 7B). Within the outer mural region, all cells contained NPR2 protein, in approximately equal amounts; the heterogeneity seen for the LHR was not evident (Fig. 7C; compare with the wildtype follicle in Fig. 7D). Within the inner mural region, the cells closest to the antrum contained more NPR2 (Fig. 7A,C), again consistent with previous reports of mRNA distribution (4,21).

**Figure 7.**
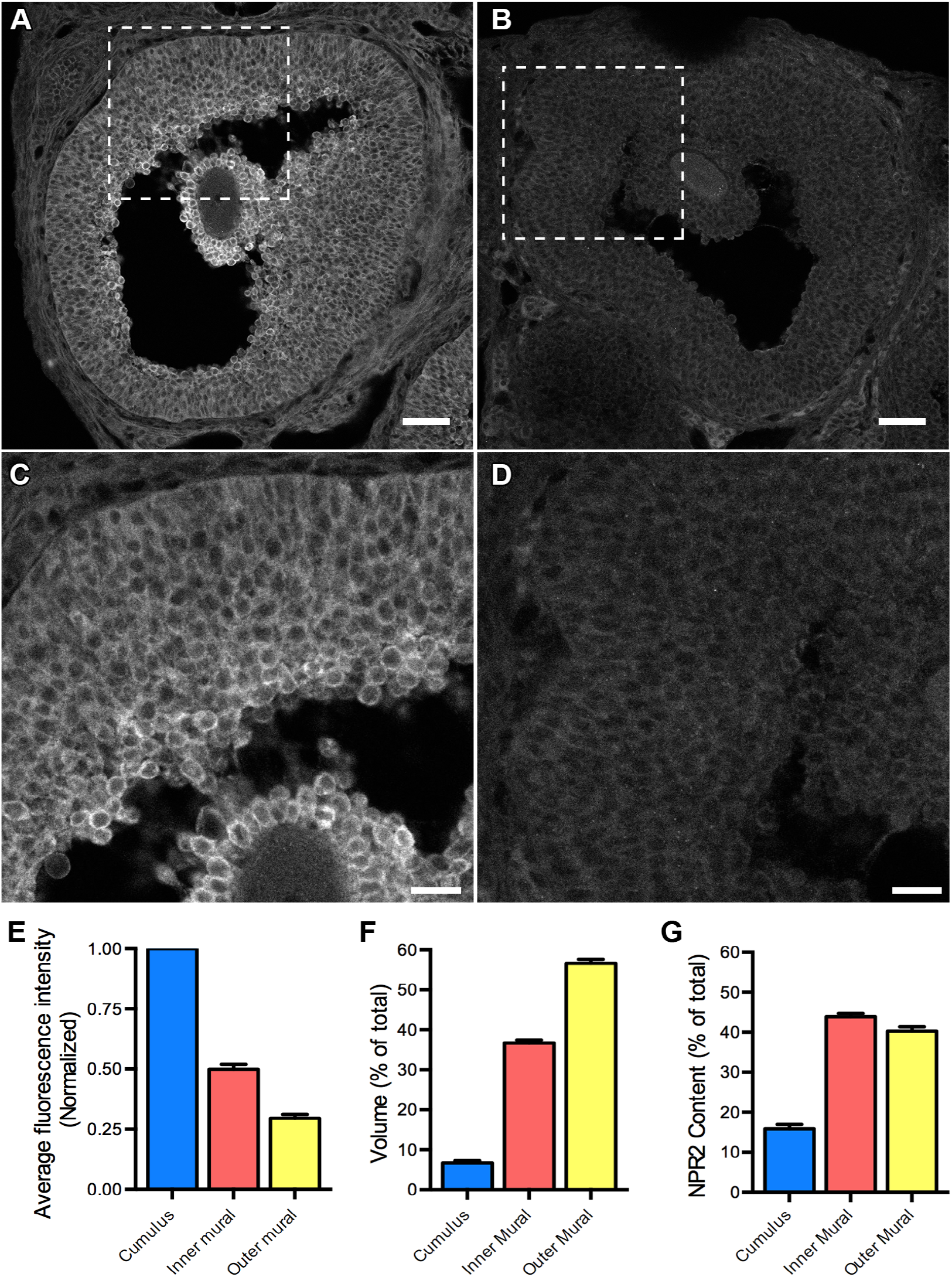
NPR2 distribution in preovulatory follicles. (A-D) Preovulatory follicles in ovaries from HA-NPR2 (A,C) or wildtype (B,D) mice. All sections were labeled with an HA tag antibody. HA-NPR2 and wildtype sections were labeled and imaged using identical conditions. Single optical sections are shown. Scale bars indicate 50 *μ*m for A and B, 20 *μ*m for C and D. (E) Average fluorescence intensities for each region of HA-NPR2 follicles, after subtraction of background determined from measurements of wildtype follicles. Values are normalized to cumulus intensity. Data are from 33 follicles from 5 HA-NPR2 mice, and 22 follicles from 4 wildtype mice. (F) Relative volumes of cumulus, inner mural, and outer mural regions, determined by counting the number of cells in each region in equatorial sections. Cell counts were used to estimate the relative volumes of the three regions, assuming volumes are proportional to the number of cells. Data are from 10 follicles from 4 mice. (G) Percentage of total NPR2 content in cumulus, inner mural, and outer mural. Values were determined by multiplying numbers from E and F and are expressed as a percentage of the total NPR2 content. Graphs show mean ± SEM.

Measurements of HA-NPR2 fluorescence intensity indicated that the average concentration of NPR2 in the outer mural region is about 30% of that in the cumulus cells (Fig. 7E). On average, the concentration of NPR2 in the inner mural region is about 50% of that in the cumulus cells (Fig. 7E). We then estimated the volume of each of the three regions, by counting the number of cell nuclei in each region of an equatorial section, and assuming that the volumes of individual cells in each region were similar. This procedure yielded the graph shown in Fig. 7F. Multiplying the values in Fig. 7E by those in Fig. 7F yielded the graph shown in Fig. 7G, which indicates that about 40% of the total NPR2 is in the outer mural cells, about 44% in the inner mural cells, and about 16% in the cumulus cells.

### Determination of the percentage of the total NPR2 that is in LHR-expressing cells

Based on the data from Fig. 7G showing that about 40% of the total NPR2 protein is in the outer mural cells, and the data from Fig. 4C showing that the percentage of outer mural cells that express the LHR is about 13-48%, we concluded that only about 5-19% of the total follicle NPR2 protein is in cells that also express the LHR (40% × 13-48%). Ideally different epitope tags would have been used for the LHR and NPR2, allowing localization of the 2 proteins in the same individual cells. However, as discussed above, only the HA epitope tag provided specific labeling.

This calculation of the percentage of the total NPR2 protein that is in LHR-expressing cells relies on our findings that essentially all of the LHR is in the outer mural cells (Figs 3,4), and that the NPR2 concentration is similar in all of the outer mural cells (Fig. 7C). Assuming that NPR2 in all of these cells is equally active, our findings indicate that to explain how LH signaling decreases NPR2 activity in the follicle to 50% of the basal level (10, 11), signaling by the LHR would need to inactivate NPR2 not only in cells that co-express the LHR, but also in neighboring cells that do not express the LHR.

### Attenuation of the LH-induced cGMP decrease in the mural granulosa cells by an inhibitor of gap junction permeability

One of the possible mechanisms that could convey the NPR2-inactivating signal from a cell expressing the LHR to a non-LHR expressing neighbor cell is diffusion of a small molecule through gap junctions. To investigate this possibility, we used mice expressing an optical cGMP sensor (cGi500) to measure the LH-induced cGMP decrease in the outer mural granulosa cells of isolated follicles, and incubated the follicles with CBX to decrease gap junction permeability.

In the presence of CBX, the LH-induced cGMP decrease in the outer mural cells was partially attenuated (Fig. 8A,B). CBX has been previously shown to effectively inhibit connexin 37 junctions between the oocyte and cumulus cells (6). Tests of gap junction permeability, using fluorescence recovery after photobleaching (FRAP), confirmed that under the conditions used, CBX completely blocked diffusion of a fluorescent tracer (Alexa 488) through the connexin 43 junctions between the mural granulosa cells (Fig. 8C,D). These findings indicate that gap junctions are a mediator of the LH-induced cGMP decrease in the outer mural granulosa cells, consistent with this mechanism contributing to inactivation of NPR2 in cells other than those expressing the LHR.

**Figure 8.**
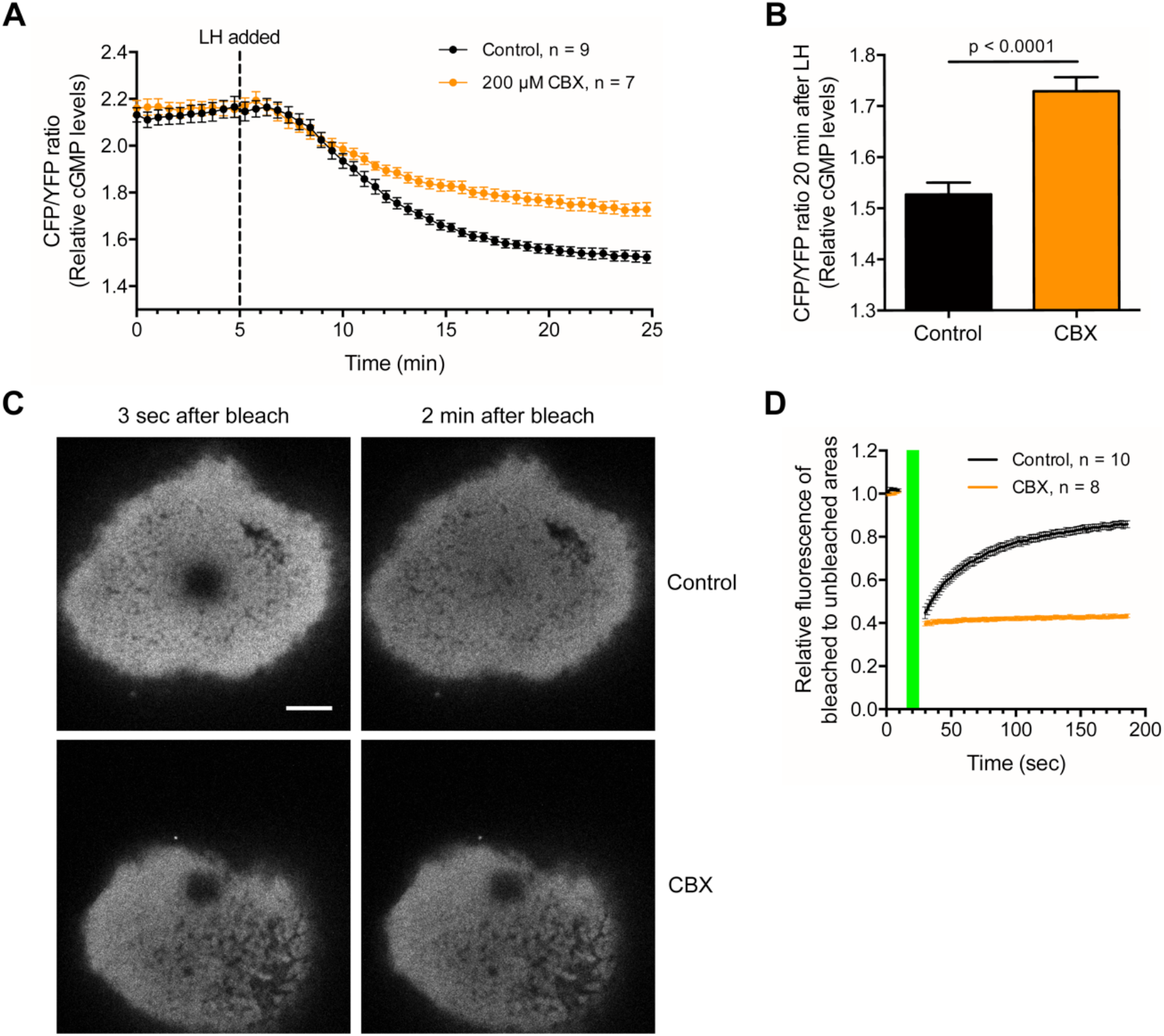
Attenuation of the LH-induced cGMP decrease in the outer mural granulosa cells by inhibition of gap junctions. (A) Follicles expressing the cGi500 FRET sensor for cGMP were preincubated for 2 hours with or without 200 *μ*M CBX and then imaged every 30 seconds by confocal microscopy, before and after addition of 10 nM LH. A decrease in cGi500 CFP/YFP emission ratio indicates a decrease in cGMP. The graph shows CFP/YFP ratios as a function of time after LH perfusion, mean ± SEM for the indicated number of follicles. (B) CFP/YFP ratios, mean ± SEM, at 20 min after perfusion of 10 nM LH, with or without CBX preincubation (data from A). The bars indicate the average of the final 5 scans of the recordings shown in A. Data were analyzed by an unpaired t-test. (C) CBX effectively blocks gap junction communication, as tested by FRAP in the outer mural granulosa cells of follicles preincubated for 2 hours with or without 200 *μ*M CBX. Alexa-488 was injected into the oocyte within the follicle and was allowed to diffuse through gap junctions into the outer granulosa cells for at least 2 hours prior to CBX treatment (14). Images show a field of outer mural granulosa cells 20 *μ*m deep into the follicle. Images on the left were collected as quickly as possible (~3 sec) after photobleaching; images on the right were obtained 2 min after photobleaching from a series of 100 scans taken every 1.6 sec after the end of photobleaching. Scale bar indicates 50 *μ*m. (D) Quantitation of FRAP in control follicles and follicles preincubated for 2 hours with 200 *μ*M CBX. To correct for absolute differences in fluorescence intensity and mild photobleaching during image collection, the fluorescence of the photobleached area was normalized to a non-bleached region of similar initial intensity. The green bar represents the period of photobleaching, with a series of 6 scans collected every 1.6 sec before bleaching and a series of 100 scans collected every 1.6 sec after bleaching. Data points indicate the mean ± SEM for 10 control and 8 CBX FRAP trials using 2 control and 2 CBX-treated follicles.

## Discussion

Immunofluorescence localization of cellular proteins that are expressed at low levels requires the availability of highly specific antibodies, as well as a control sample lacking the antigen. For studies of endogenous proteins in tissues, these requirements can be met by use of mice in which the protein of interest is replaced with an epitope-tagged protein for which specific antibodies to the tag are available (23,24). Wildtype mice lacking the tag provide a negative control. Here we produced mice in which the LHR and the NPR2 guanylyl cyclase were replaced with HA-tagged proteins.

Using these mice, we quantified LHR and NPR2 protein localization in ovarian follicles. Our results are consistent with mRNA and ligand binding localization seen in previous studies, but provide cellular vs tissue level resolution. The results showed that only some of the outer mural granulosa cells express the LHR. Images in previous studies also suggest heterogeneity in LHR distribution in different regions of the outer mural granulosa cells (15–20), but the resolution of these earlier methods was insufficient to detect cellular heterogeneity. By quantifying the distribution of the LHR and NPR2 in different regions of the follicle, we determined that the cells that express the LHR contain less than 20% of the total NPR2 in the follicle.

Our results raise the question of how LHR signaling in a subset of the granulosa cells that contains less than 20% of the total NPR2 protein can account for the measured decrease in NPR2 activity to a level of ~50% of that before LH application (10,11). Our findings support a role for intercellular diffusion of a small molecule through gap junctions in conveying the NPR2-inactivating signal to mural granulosa and cumulus cells that do not express the LH receptor. Other mechanisms may contribute as well, including the release of epidermal growth factor receptor (EGFR) ligands that diffuse extracellularly within the follicle (1,47). Inhibiting EGFR kinase activity attenuates the LH-induced cGMP decrease to varying degrees with different experimental conditions (1,47), but measurements with the cGi500 FRET sensor showed that preventing EGFR activation has little or no effect on the initial LH-induced decrease in cGMP (14).

A candidate for the small molecule that is generated by LH signaling and diffuses through gap junctions to inactivate NPR2 in adjacent cells is cAMP. A role for cAMP is supported by evidence that elevating cAMP with forskolin mimics the LH-induced cGMP decrease in the follicle (14). Furthermore, LH signaling, via the cAMP-activated protein kinase A, phosphorylates and activates a regulatory subunit of the PPP2 phosphatase in rat granulosa cells (48), and PPP family phosphatase activity is required for NPR2 dephosphorylation in response to LH (11). Our finding that CBX attenuates the LH-induced cGMP decrease in the outer mural granulosa cells is consistent with this model.

Our findings, as well as previous studies of the localization of the LHR and NPR2 (see Introduction) also raise the question of what factors determine the localized expression of these proteins within the tissue. Both stimulatory and inhibitory factors have been identified. One essential factor that stimulates LHR expression is FSH (15,16,41; see Fig. 2C). Although FSH receptors are present throughout the mural and cumulus granulosa cells (18), FSH delivery to the follicle from blood vessels outside of the follicles results in a gradient of FSH within the follicle, with more FSH near the basal lamina (16). This gradient of FSH could contribute to the observed gradient of LHR expression. It is unknown whether FSH receptor expression in the granulosa cells might be heterogeneous on a cellular level. If so, this could contribute to the heterogeneity of LHR expression.

Contact with the extracellular matrix of the basal lamina (41, 49) is another factor that stimulates LHR expression. Our findings indicate that this contact is not sufficient for LHR expression. Androgens that are produced by surrounding theca cells also stimulate LHR expression (50, 51), and this may be an additional factor that influences the localization of the LHR.

In contrast to these positive factors, oocyte-secreted proteins of the TGF-beta superfamily, growth differentiation factor 9 and bone morphogenetic protein 15, inhibit LHR expression (41, 52). These same oocyte-secreted proteins that inhibit LHR expression stimulate NPR2 expression (4), supporting the concept that oocyte-secreted proteins are a cause of higher NPR2 expression and lower LHR expression in granulosa cells nearer to the oocyte. Long cellular processes that extend from the outer granulosa cells (37) could potentially sense a concentration gradient of TGF-beta proteins across the mural granulosa cell layer. In future studies, it will be of interest to investigate the spatial distribution of this gradient.

In future studies of the function of the LHR and NPR2 in ovarian granulosa cells, the HA-tagged LHR and NPR2 mice described here should be useful for determining whether the localization of these proteins changes in response to LH signaling. Previous studies of cultured cells exogenously expressing the LHR indicated that LH binding results in internalization of the receptor, thus prolonging and diversifying the hormonal signal (53, 54). However, imaging of LH-induced internalization of the LHR in intact ovaries has not been accomplished. The ability to directly visualize LHR and NPR2 proteins in the ovarian follicle may also facilitate investigation of regulatory factors that determine their localized expression.

In addition to its function in the granulosa cells, the LHR regulates steroid production by ovarian cells outside of the follicle itself (theca/interstitial cells) (55) and by testicular Leydig cells (56). The LHR is also expressed in the corpus luteum, although its contribution to stimulating progesterone production to support pregnancy in mice is not clearly understood (57). Outside of the ovary, the LHR is expressed in the uterus and oviduct (58), although LHR expression in these tissues is not required for reproduction (59). Outside of the reproductive system, recent evidence indicates that the LHR functions in hematopoietic stem cells (60). Likewise the NPR2 guanylyl cyclase functions in the skeletal, cardiovascular, and nervous systems (61), but its localization is not well characterized. As we have found for ovarian granulosa cells, more precise knowledge of LHR and NPR2 localization obtained using mice with HA-tagged versions of these proteins may contribute to understanding of their function in cell signaling in diverse physiological systems.

## Acknowledgments

We thank Deborah Kaback for her help in generating and maintaining the genetically modified mice, and Robert Feil for providing the cyclic GMP sensor mice. We also thank Luisa Lestz, Leia Shuhaibar, and Giulia Vigone for their assistance with experiments, and Daniel Bernard, Raj Duggavathi, John Eppig, Aylin Hanyaloglu, Ilpo Huhtaniemi, Eric Levine, Olga Morozova, Bruce Murphy, Rachael Norris, Lincoln Potter, JoAnne Richards, Adolfo Rivera-Müller, Alexander Sorkin, and Stephen Yeung for advice and helpful discussions.

## Legends for supplementary videos

**Video 1.** A series of confocal sections taken at 1 *μ*m intervals through a 10 *μ*m thick physical section of a preovulatory follicle from an ovary of an immature HA-LHR mouse that was injected 44 hours previously with eCG. HA-LHR in green, DAPI-labeled nuclei in blue. Scale bar = 10 *μ*m.

**Video 2.** A series of 100 serial sections, imaged by scanning electron microscopy, of outer mural granulosa cells of a preovulatory follicle. Each section is 30 *μ*m × 30 *μ*m × 65 nm. Figures 5C-E show images and reconstructions made from a larger volume from which this stack was cropped. Scale bar = 3 *μ*m.

**Video 3.** Reconstruction of 9 outer mural granulosa cells from a 70 *μ*m × 70 *μ*m × 14 *μ*m volume of serial sections, imaged by scanning electron microscopy. LHR-expressing cells are shown in shades of green and non-LHR-expressing cells shown in shades of blue.

**Video 4.** Reconstruction of 2 outer mural granulosa cells selected from the reconstruction in video 3, showing gap junctions between these 2 cells in orange and gap junctions with other neighboring cells in yellow. The LHR-expressing cell (green) was sliced to show gap junctions between the two cells.

